# A macroecological description of alternative stable states reproduces intra- and inter-host variability of gut microbiome

**DOI:** 10.1101/2021.02.12.430897

**Authors:** Silvia Zaoli, Jacopo Grilli

## Abstract

The most fundamental questions in microbial ecology concern the diversity and variability of communities. Their composition varies widely across space and time, as it is determined by a non-trivial combination of stochastic and deterministic processes. The interplay between non-linear community dynamics and environmental fluctuations determines the rich statistical structure of community variability, with both rapid temporal dynamics fluctuations and non-trivial correlations across habitats. Here we analyze long time-series of gut microbiome and compare intra- and inter-community dissimilarity. Under a macroecological framework we characterize their statistical properties. We show that most taxa have large but stationary fluctuations over time, while a minority is characterized by quick changes of average abundance which cluster in time, suggesting the presence of alternative stable states. We disentangle inter-individual variability in a major stochastic component and a deterministic one, the latter recapitulated by differences in the carrying capacities of taxa. Finally, we develop a model which includes environmental fluctuations and alternative stable states. This model quantitatively predicts the statistical properties of both intra- and inter-individual community variability, therefore summarizing variation in a unique macroecological framework.

---

Microbial communities are the prototypical complex, high-dimensional, ecosystems. Their diversity is astonishing and occurs at all scales, with many taxa in the same communities and many strains coexisting within each taxon. Their dynamics is driven by mechanisms operating at all temporal scales [1, 2]: gut microbiota composition change in response to diet or other external stimuli on a daily basis [3], and with aging through the course of a lifetime [4, 5]. Similarly, multiple processes determine the variation in composition across hosts, ranging from behavioral [6] to genetic factors [7].

This high-dimensional complex space of variation in community composition might conceal low-dimensional structures. For instance, it has been proposed that community compositions, when compared across hosts, cluster in a few discrete enterotypes characterized by specific taxa [8, 9] as the result of life-history characteristics. These clusters are thought to correspond to different functional properties, for instance summarized in the enrichment of different metabolic pathways [8]. Their existence and biological significance are not broadly accepted. There is in fact evidence that inter-host variation of gut community composition is continuous, with the existence of discrete types being only apparent as a result of statistical artifacts [10].

The existence of discrete clusters in community composition also concerns intra-host variability. In fact, the fluctuating composition of a host’s community might also cluster around several alternative compositions. The long autocorrelation times make it particularly challenging to formulate robust statistical methods to infer enterotypes [11].

Whether variation is discrete or continuous has important practical consequences. For instance, if cluster exists they could provide a biomarker of health-conditions and target for clinical purposes [9, 10]. Beyond the health-related applications, there are important conceptual consequences. The existence of enterotypes would be an evidence that the dynamics of gut microbial communities results in alternative stables states, which appear as consequence of the interplay between dynamics and environmental forcing.

Understanding mechanistically the properties of microbial communities is especially difficult, as the complexity of realistic mechanistic models cannot be simply related to the structure of their dynamical attractors. This represents one of the main challenge to understand quantitatively and mechanistically the properties of microbial communities.

Macroecology, which focuses on the statistical patterns of community composition, offers a promising strategy to characterize quantitatively microbial communities. Recent works have characterized the macroecological patterns of community composition [12, 13]. Under the macroecological lens, community dynamics is characterized by large and rapid fluctuations with robust and reproducible statistical properties [14–16]. These regular patterns are well described by the stochastic logistic model with environmental noise, which assumes that fast environmental fluctuations perturb abundances around a taxa-specific carrying capacity [15, 16].

Here we study the intra-host dynamics and inter-host variability under a unique macroecological framework using the time series of 14 individuals. We show that most of the taxa display stationary dynamics [11], with rapid fluctuations around a constant carrying capacity, as predicted by the stochastic logistic model. The non-stable OTUs are characterized by discrete shifts of the carrying capacity, which occur coordinately in time for several taxa. This suggests that the change of community composition in time proceeds by discrete changes of the carrying capacities. Surprisingly, differences in carrying capacity are also what explains the variability across hosts, in combination with independent stochastic fluctuations. We finally show that both the intra-host and inter-host variability can be quantitatively captured by a stochastic logistic model modified to include the existence of alternative stable states.

## I. RESULTS

### A. Characterizing long-term community dynamics with dissimilarity

The relative abundances of OTUs within a community have wide fluctuations in time (see Fig 1A). In this context, we may ask whether the difference in community composition in two snapshots of the same community at different times are only due to wide but stationary fluctuations or if, instead, the composition changes in time. In the first scenario, abundances would fluctuate around constant averages, with constant amplitudes (Fig. 1B). In the second scenario, average abundances would change in time, either continuously or discretely (Fig. 1C), possibly due to jumps between alternative stable states. We can discriminate between the two scenarios by looking at a dissimilarity measure quantifying how the abundance of each OTU becomes different from itself at a certain time lag. Let us call this dissimilarity Φ(*T*). In the first scenario (Fig. 1D), Φ(*T*) for each OTU would show an initial increase, over the short time-scale of relaxation to stationarity, and then would reach a plateau at the level of self-dissimilarity of a fluctuating random variable, which depends on the amplitude of the fluctuations. In the second scenario (Fig. 1E), instead, the dissimilarity would continue to increase over longer time-scales due to the abundance getting more and more different from its past self.

**FIG. 1:**
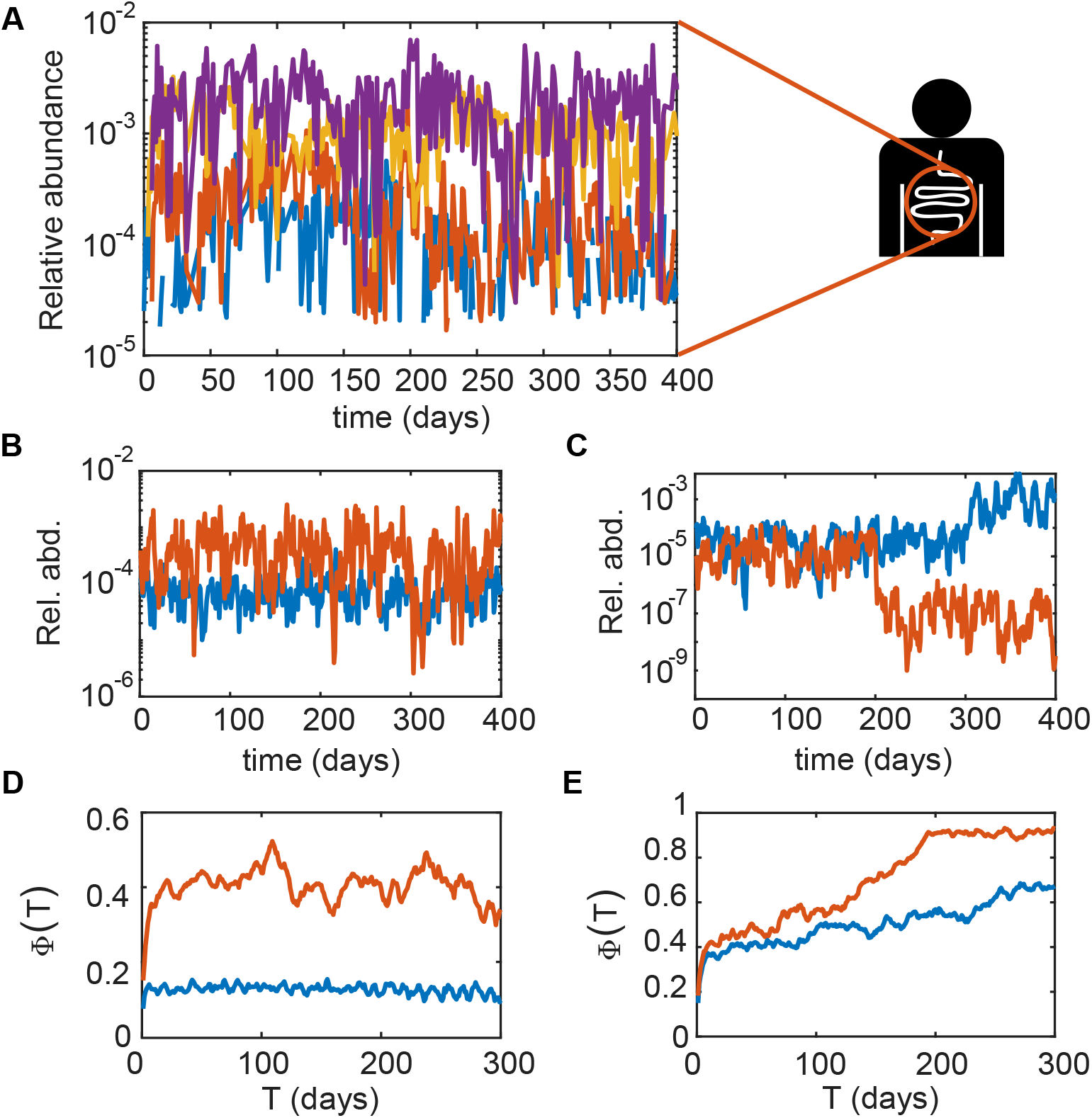
Characterizing long-term community dynamics with dissimilarity. A) Examples of relative abundance trajectories with wide short-term fluctuations; B) Relative abundances trajectories with carrying capacity and noise intensity constant in time; C) Relative abundances trajectories with carrying capacity changing in time and constant noise intensity. D) Dissimilarity Φ(*T*) for varying lag *T* for the abundance trajectories in panel B; E) Dissimilarity Φ(*T*) for varying lag *T* for the abundance trajectories in panel C.

### B. Within-individual dissimilarity displays two long-term behaviors

We analyzed the time-series of of 14 individuals’ gut microbial communities from three different datasets (see Methods). In each time-series, we compute the dissimilarity Φo_*i*_(*T*) for each OTU according to a dissimilarity measure that is not subject to the bias introduced by random sampling of the community (see Methods). Examples of curves Φ_*i*_(*T*) obtained for some OTUs are plotted in Figure 2A. Both the short-time and long-time behaviors appear to differ widely between OTUs. In order to asses the significance of these behaviors we compare them with what is expected from a null model.

**FIG. 2:**
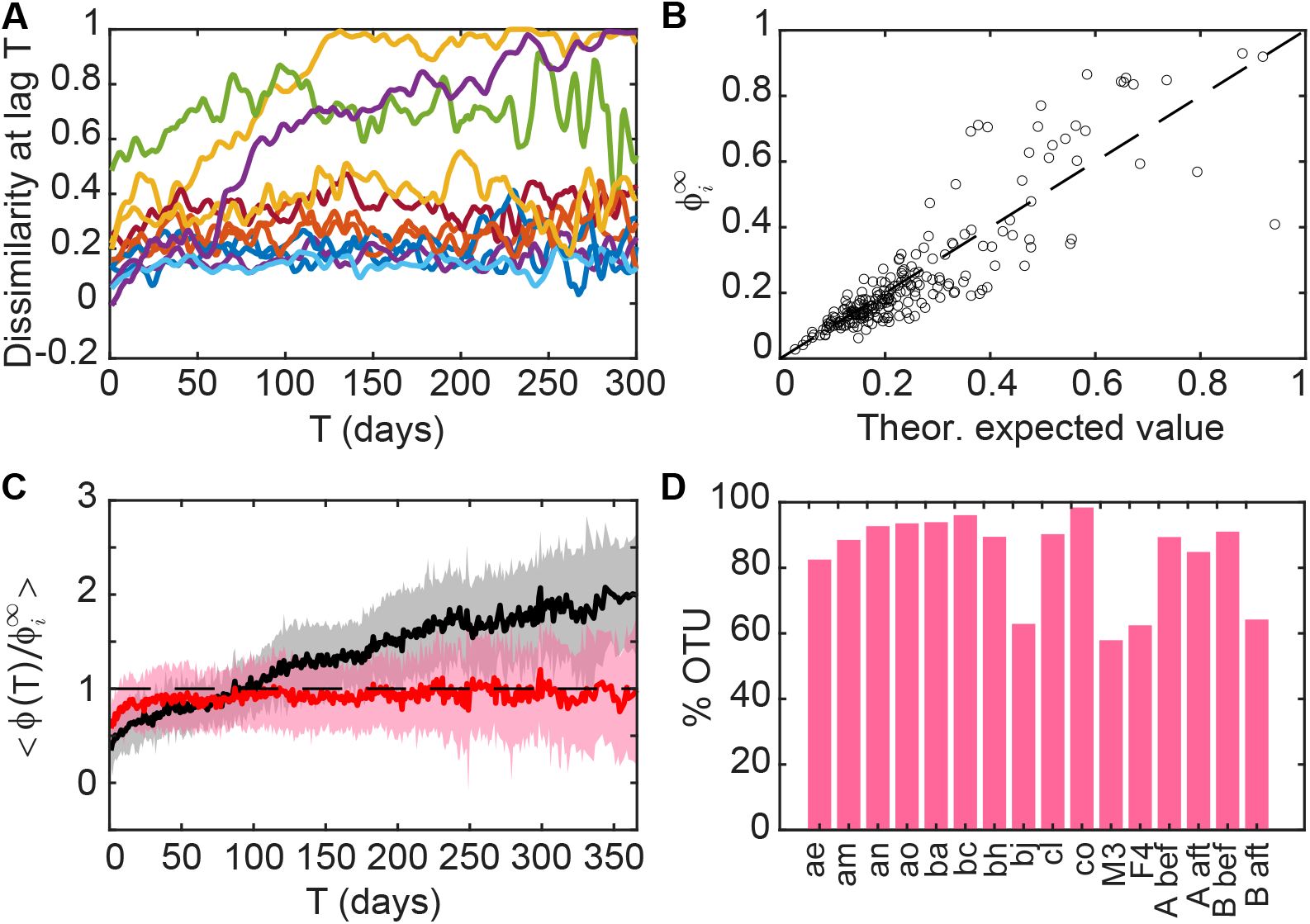
Dissimilarity indicates that most OTUs have stationary dynamics, but a minority has non-stationary behaviour. A) Examples of Φ_*i*_(*T*) curves of individual OTUs; B) Scatter plot of the theoretical prediction for the asymptotic value of Φ_i_(*T*) for an OTU with noise intensity *σ* against 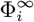, the empirical average value of Φ_i_(*T*) for *T* > 10 days, for individual ‘bh’. The 1:1 line is plotted as reference; C) Average of Φ(*T*)/Φ^∞^ over OTUs where it is classified as flat (red curve) and increasing (black curve), for individual ‘bh’. Shaded areas represent one standard deviation intervals; D) Percentage of OTU whose Φ(*T*)/Φ^∞^ is classified as flat in each individual.

The Stochastic Logistic Model with environmental noise (SLM, see Methods) is known to describe several statistical properties of abundance fluctuations for microbial species [15, 16]. In particular, it predicts that the distribution of fluctuations at stationarity is a Gamma distribution, which depends on two parameters: *K_i_*, the carrying capacity appearing in the SLM, and *σ_i_*, which can be interpreted as the amplitude of environmental noise. These two parameters depend on the identity of OTU *i*. The noise *σ_i_* can be expressed in terms of the coefficient of variation of abundances. As a consequence, Taylor’s law on abundance fluctuation with exponent 2 [16] implies that *σ_i_* and *K_i_* are not correlated across OTUs.

Assuming that the abundance dynamics follows the SLM with parameters that remain constant in time, Φ_i_(*T*) has an initial increase and then reaches a plateau. The height of this plateau can be computed analytically as the expected value of Φ_i_(*T*) for large *T* by taking the abundances of the OTU at lag *T* ≫ 1 as independent Gamma random variables with the same parameters (see Methods). This analytical expectation depends indeed only on the parameter *σ_i_*, which can be directly estimated from the data (see Methods). Fig. 2B shows that this expected value 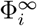 correlates well across OTUs with the empirical height of the plateau, measured as the mean of Φ_i_(*T*) over all *T* > 10 days.

According to the SLM, when normalized by 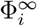, all the individual curves Φ_*i*_(*T*) should have a plateau at 1. Therefore, if the empirical normalized dissimilarity curve of an OTU has a plateau at 1, it means that for that OTU the level of dissimilarity at large lags is explained only by the stationary fluctuations of abundance. For such an OTU we can conclude that the abundance dynamics has constant parameters over the observed time interval (ranging from 6 months to 1.5 y for the analyzed individuals). This behavior is what we observe for the majority of OTUs in each individual.

We classified the normalized dissimilarity curves 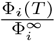 as either ‘flat’ (if they plateau after the initial transient) or ‘increasing’ (if they keep increasing, see Methods), and computed the average curves within the two categories (Fig. 2C). The average of the flat curves (red line) reaches a plateau of 1, indicating that the OTUs falling in this group have a stationary abundance dynamics. These stable OTUs are the majority, representing on average 84% of the total in each individual (Fig. 2D). Other OTUs do not reach a plateau, and their normalized dissimilarity grows over all the observed time interval (black line in Fig. 2C).

These results show that a large part of the OTUs in a gut microbial community fluctuates around a stable average composition on a ~1 y timescale, while a smaller fraction has a non-stationary behavior. We explore more carefully the properties of non-stationary OTUs in the next section.

### C. Not-stationary OTUs are characterized by transitions between alternative values of the carrying capacity

If the abundance evolves according to the SLM with constant parameters, the dissimilarity Φ(*T*) reaches a plateau Φ_∞_ determined by *σ*. The dynamics of non-stationary OTUs, for which we observe an increasing Φ(*T*), could potentially still be described by the SLM with time-dependent parameters *K_i_*(*T*) and *σ_i_*(*T*), which change on the time-scales of months.

Two paradigmatic behaviors could describe the time-dependency of parameters. The first is a slow variation of the parameters, where *K_i_*(*T*) and/or *σ_i_*(*T*) change steadily over time. The second one corresponds to rapid transitions to different values of parameters spaced out by time windows where parameters are constant. These represents two extreme case, and a mixture of the two could also describe the time-dependency of parameters.

A visual inspection of the abundance time-series of non-stable OTUs shows that in many cases there is a sudden jump in the relative abundance (see Figure S5 A and B) which can be interpreted as a jump in the carrying capacity *K*, suggesting that the second scenario might apply.

We introduced therefore a method, based on a quantity related to the Kullback-Leibler divergence, to detect jumps of *K* in a noisy time-series (see Methods). By applying this method we can identify, for each individual, which OTUs present jumps in their carrying capacity *K*. On average, 54% of OTUs with increasing slope display sudden jumps in their carrying capacity *K*. For 10 out of 16 time-series OTUs with jumps of *K* are over-represented among OTUs with increasing Φ_*i*_(*T*) (p-value<0.05 according to a hypergeometric test, see Fig. S5 C and Table S1). Cases where Φ_*i*_(*T*) is increasing but no jump of *K* is detected can be due to noise in the estimate of Φ_*i*_(*T*) or to jumps falling below the detection threshold of our method. There are also cases where a jump in *K* is detected but Φ_*i*_(*T*) is flat. These are either cases where the change in *K* is short-lived or where the jump is small. Changes in *K* would in fact affect the value of Φ_*i*_(*T*) only is they are detectable with respect to the every-day abundance fluctuations. Examples of all these cases are shown in Fig. S7.

The majority of OTUs displaying non-stationary dynamics are therefore associated to rapid jumps between alternative values of *K*. Interestingly, these sudden jumps are not randomly distributed over the time-span of the time-series. We observe in fact that, for individuals where several OTUs have jumps in *K*, the times where transitions happen are localized at specific times (see Fig. S8). This clustering of transitions strongly suggests that jumps in K are driven by external conditions which affect multiple OTUs at the same time.

The dynamics of OTUs abundance within an individual is therefore characterized by two main timescales. A short-timescale corresponding to rapid fluctuations, which are described by the noise term appearing in the SLM, and a larger time-scale which characterizes the frequency of the transition between alternative values of *K*.

### D. Dissimilarity between individuals is quantitatively reproduced by a combination of stochastic fluctuations and different carrying capacities

Within an individual, we found that similarity is determined by rapid fluctuations around a constant composition and by occasional sudden jumps of the carrying capacity. It is well known that some features of microbiome composition remains stable for years and are host-specific. In this context, we now investigate the dissimilarity between individuals, in order to quantify the role of different sources of variation. In particular we aim at disentangling the contribution of stochastic effects that are independent in the two individuals from that of reproducible differences between them. Under our framework, the stochastic aspects correspond to the rapid fluctuations of abundance, which are statistically independent between hosts. The deterministic factors correspond instead to different parameters *K* and *σ* that characterize the dynamics of each OTUs in different hosts.

In order to quantify the importance of the two causes of dissimilarity, we compare the empirical dissimilarity with that predicted by the SLM under different hypotheses on the sources of variation that are present. For each OTU i we compute the empirical dissimilarity between two individual *a* and *b* averaged over all lags *T*, 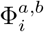. We then consider a first null model that includes both sources of variation: the dynamics of OTU i is described by the SLM with different parameters in the two individuals *a* and *b*, 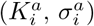 and 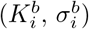, which are estimated using the full-time series (see Methods). The difference in these parameters between the individuals aim at capturing the host-specific factors that determine community composition. The rapid fluctuations are instead determined by the stochastic term in the SLM. In the Methods, we obtain an analytical prediction for the dissimilarity 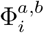, which matches the empirical value with high accuracy (Fig. 3A and Fig.S9). This result implies that the host-specific aspects of community composition can be effectively captured in the variability of the parameters *K* and *σ*.

**FIG. 3:**
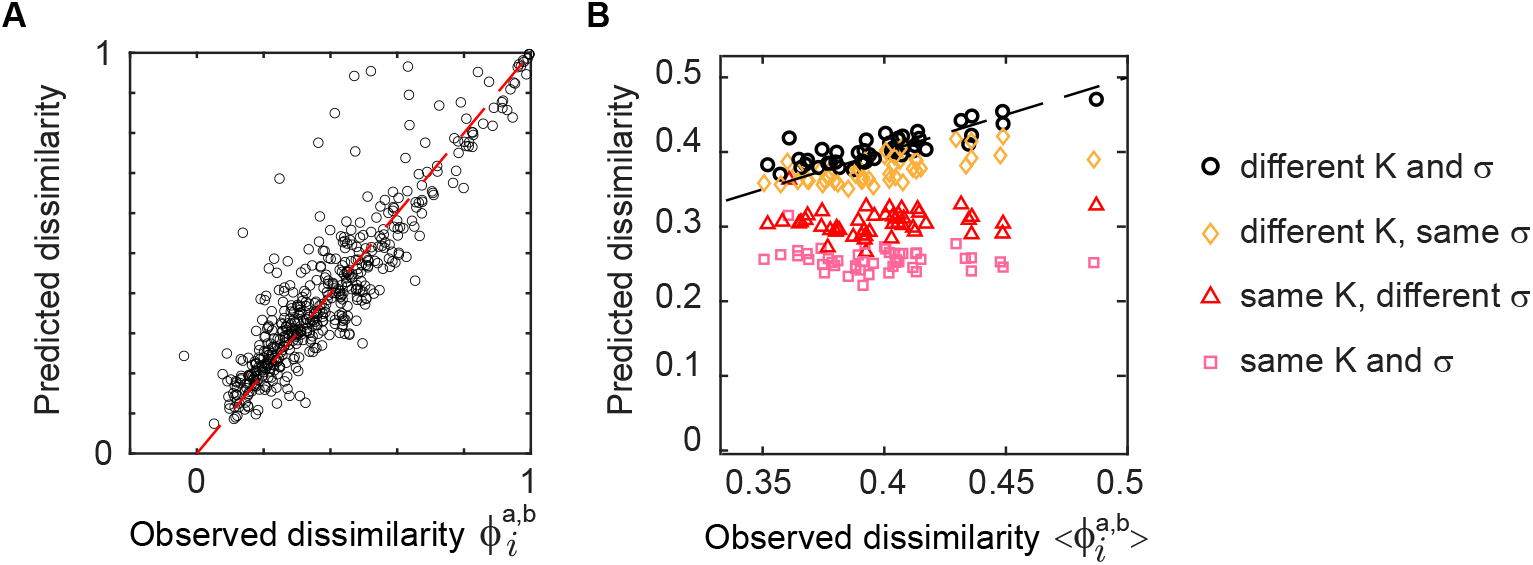
Dissimilarity between individuals is explained by independent stochastic fluctuations and differences in carrying capacities. A) Comparison of the observed dissimilarity 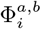 of OTU *i* between the two individuals ‘am’ and ‘bh’ with its theoretical expected value computed with the individual parameters estimated for each individual. Each point represents an OTU; B) Comparison between the average over OTUs of the empirical dissimilarity between individuals *a* and *b*, 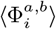, with its theoretical expected value computed using either the individual parameters estimated for each individual or the same parameters for both individuals (see legend). In the latter case, the parameters are set to the average of the individuals’ parameters. Each point corresponds to a pair of individuals from the same dataset. The 1:1 line is shown as reference.

To assess how much of the dissimilarity is due to the difference in *K* and *σ* between the two individuals and how much to the stochastic abundance fluctuations, we consider a hierarchy of four null models, which differ in whether *K* and/or *σ* vary between hosts. Fig. 3B compares the predictions of these four null models with the dissimilarity 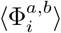 averaged across OTUs.

The simplest scenario of the four is the one where the predicted dissimilarity between individuals is only due to stochastic effects, i.e. *K* and *σ* do not differ between individuals. Fig. 3B shows that stochastic effects alone — as expected — do not fully explain the dissimilarity between individuals. They are nevertheless the main contributor to the variability between individuals, capturing on average 72% of it (see also Fig. S10).

Consistent with what shown in Fig. 3A for individual OTUs, Fig. 3B also shows that considering the empirical variability of both *K* and *σ* across hosts fully captures the empirical dissimilarity. On the other hand, *K* and *σ* do not contribute equally to dissimilarity. Considering only the variation of *σ*, while keeping *K* constant across hosts, performs only slightly better than the model with only stochasticity, as it explain on average 84% of the dissimilarity. On the opposite, keeping *σ* fixed, while *K* is allowed to vary across host predicts much better the observed dissimilarity, with an average 96% accuracy.

These results imply that the difference in community composition between individuals can be explained by considering the combination of independent stochastic effects, as modeled by the SLM, and deterministic factors. The latter can be captured by differences the carrying capacity *K_i_* of OTUs in the two hosts.

### E. The SLM with alternative stable states reproduces inter- and intra-host variability

The parallel between the long-time dynamics within an individual, characterized by discrete transitions of the carrying capacity *K*, and the difference between individuals, whose deterministic part is largely determined by differences in the carrying capacity, suggests the possibility to formulate a dynamical model capturing both.

We assume that each OTU has a characteristic, host-independent, value of the carrying capacity, 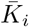. Within an individual, the actual carrying capacity of an OTU at a given time is 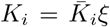 where *ξ* is a Lognormal random variable with average 1. The value of the carrying capacity is kept constant over time, until a transition happens to a new value 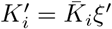 obtained by drawing a new value of the random variable *ξ*. Similarly, the values of the carrying capacity in two different individuals are obtained with the same model by considering independent realizations of the random variable *ξ*. This model aims at capturing the statistical properties of the values of *K_i_*, within and across hosts. The strong assumption is that the fluctuations within and across hosts can be captured under the same framework.

The free parameters of this model are the variances of the random variable *ξ*, which determine how much *K_i_* varies over time and across hosts. In the Supplementary Information, we estimate empirically the values of the variance of *ξ* across hosts for each individual OTU. The results show that the variances do not differ much across OTUs, and are statistically compatible (see Fig. S4). We therefore choose the variance of to be OTU independent, obtaining a model with no free parameter, as both the 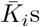 and the variance of *ξ* can be estimated directly from the data.

To test if this model can indeed reproduce the statistical properties of the dynamics within an individual and the differences between individuals, we used it to simulate the abundance time-series of each individual and compared the results with those obtained from the empirical time-series.

We simulated the abundance time-series of each individual according to a SLM with carrying capacity that jumps according to a Poisson process and assumes new values according to the model described above (see Methods). Repeating on these simulated data the analysis performed on empirical data, we find that the intra-individual dissimilarity behaves very similarly to the empirical one (Fig. 4A). Moreover, by fixing for each individual the average number of OTUs for which a jump happens in the observation window according to the empirical observations, the percentages of OTUs with increasing dissimilarity are similar to the empirical ones (see Fig. S12).

**FIG. 4:**
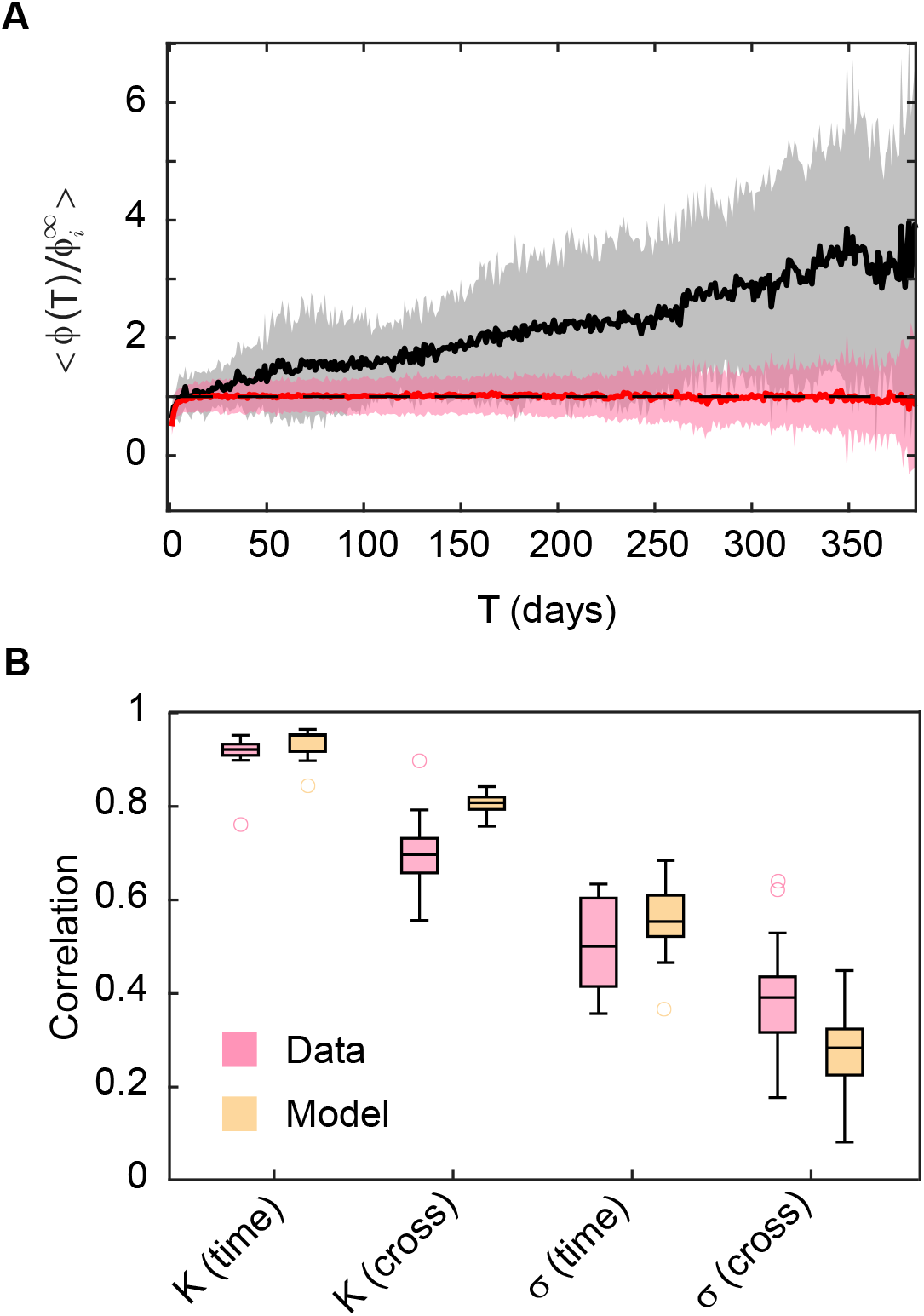
A model with stochastic fluctuations and discrete jumps of the carrying capacity reproduces the statistical properties of the dynamics within one individual and the difference between individuals. A) Equivalent of Fig. 2C obtained from time-series simulated according to a SLM model with jumps in *K* (see text for details); B) Correlation of the parameters *K* and *σ* in time (between two halves of a time series) and between pairs of individuals from the same dataset, according to the data and to the simulated time-series. For K, which ranges over several orders of magnitude, we compute the correlations of log(*K*) to avoid the correlation being dominated by a few points with large *K*.

The analysis of dissimilarity between hosts revealed that it can be fully captured by considering the difference in the values of *K* and *σ*. We therefore consider the correlation of these two parameters within hosts (for two separate windows of time) and across hosts as a relevant statistical feature of the dynamics that our model should reproduce. Fig. 4B reports the empirical values of the correlations (see also Fig. S13). In agreement with what we found previously, the carrying capacities are more correlated within host than between, indicating that their values contribute significantly to the dissimilarity between individuals. On the other hand, the values of *σ* are much less correlated within individuals and their correlation do not differ greatly between and within hosts, in agreement with the observation that variability in *σ* does not contribute much to dissimilarity across individuals. Therefore, these correlations fully summarize the results of Fig. 3B.

The model accurately predicts all the correlations, both within and between individuals. Interestingly the model also captures the lower correlations between the noise intensities *σ*, even if it does not include variability of *σ* as an explicit ingredient, implying that differences in *σ*, both between individuals and within, are simply due to noise in their statistical estimates. The model is therefore able to fully capture both the variability of individuals over long periods of time as well as differences across individuals.

## II. DISCUSSION

We considered long time-series of gut microbiome composition of several individuals and studied the similarity in composition within and between hosts. Importantly, we are able to accurately predict, under a unique quantitative modeling framework, the statistical properties of the similarities both within and between individuals.

Previous studies analyzing intra- and inter-individual gut community compositions consistently found stability in time on timescales of months or years and a higher similarity within an individuals with respect to across individuals [11, 17–20]. These studies used several similarity measures to compare communities, e.g. Jaccard index, Bray-Curtis dissimilarity or Pearson correlation. Other studies used auto-correlations [11] or mean square displacement [14] of abundance time-series to investigate the long-term behavior of single OTUs. The dissimilarity measure we use is akin to the latter approach, in that it evaluates how dissimilar a single abundance time-series becomes from its past values. However, it offers two advantages. First, it corrects for sampling bias, disentangling the dissimilarity due to different community compositions from that due to random sampling, an issue often overlooked. Secondly, we can make exact analytical predictions of its long-term behavior and value according to a well-supported model of the abundance dynamics. This allows us to distinguish OTUs that are stationary from those that are not, and to dissect the inter-individual dissimilarity in its different contributions.

Within an individual, we find that the majority of OTUs is characterized by statistically stable dynamics, in agreement with previous findings [11]. Abundances fluctuate around a value constant in time, with constant amplitude. For these OTUs, the long-term dissimilarity is due to the rapid stochastic fluctuations, and its value is determined by the amplitude of such fluctuations. The Stochastic Logistic model with constant parameters predicts correctly the time dependence and the asymptotic value of dissimilarity for these OTUs based on the noise intensity σ estimated from the time-series. A consistent minority of OTUs has a dynamics characterized by processes happening on two distinct times-scales. Beyond the wide daily fluctuations, they have large abrupt transitions in their abundance. Their dynamics is therefore non-stationary, and their dissimilarity increases in time beyond the value predicted for a stationary time-series. These observations show that both the amplitude of the fluctuations of each OTU and the proportion of non-stationary OTUs contribute to determine the overall dissimilarity of an individual’s gut community over time, which we can think of as the average of the dissimilarities of all OTUs. The overall dissimilarity is therefore a complex quantity to interpret, while our OTU-based approach provides a more detailed understanding of the underlying dynamics.

The large transitions observed in non-stationary OTUs can be effectively captured in changes of the typical abundance around which they are fluctuating. In the Stochastic Logistic Model framework, these transitions correspond to a change in the value of the carrying capacity. The non-stationary aspects of OTU dynamics can therefore be fully described by switches between alternative values of the carrying capacity. By comparing which OTUs change carrying capacity across individuals, it appears that there is no taxonomic signal, suggesting that OTUs have similar likelihood of jumping between states.

Moreover, the timing of the jumps between alternative values is strongly correlated across OTUs, with windows of times where the carrying capacity of several OTUs switches and others where the dynamics is overall stationary. The natural explanation to these clustered rapid transitions is existence of alternative stable states, which differ in the carrying capacities of a small sub-set of OTUs. They could be intrinsic to the dynamics, and driven by stochasticity, or could correspond to important changes in the environmental conditions (e.g. travel or change of diet) with big impact on community composition. Our approach is explicitly agnostic on the mechanism behind the alternative stable state, revealing instead their regular statistical features.

The SLM we used to describe the dynamics comprises an auto-regressive and a non-autoregressive term. The importance of these two components in gut microbial dynamics is quite debated [11, 21]. In the context of the SLM, the parameter σ measures the relative importance of the non-autoregressive part of the dynamics. Estimates of this parameter, corrected for the sampling bias, show that it is heterogeneous across OTUs (Fig. S1) and independent of the carrying capacity [16].

The comparison between individuals revealed that most of their dissimilarity is due to the independent stochastic fluctuations of abundances. The remaining dissimilarity, due to deterministic differences between individuals, is almost completely explained by differences in the carrying capacities of OTUs. The emerging picture indicates that the gut communities of different individuals fluctuate around states with compositions that are correlated (carrying capacities have a moderate to high correlation across individuals) but different.

The observation that both the changes within an individual on the timescale of months and the differences between individuals are explained by differences in the carrying capacities motivated our model, joining the dynamics of the SLM with transitions in the carrying capacity. The model predicts quantitatively, under a unique framework, the statistical properties of within-individual dynamics and between-individuals differences. These results are consistent with the existence of a universal dynamics of gut microbiome [22], where differences between individuals are uniquely determined by which dynamical alternative stable state they occupy. Such a universal dynamics does in fact necessarily create a parallel between within-host dynamics and between-host properties, which we unveiled under our modeling framework.

Having formulated a comprehensive framework allows to summarize OTU dynamics in few parameters, with an underlying modeling interpretation. This allows to disentangle, both conceptually and practically, two very different sources of variation: the rapid stochasticity and the jumps between alternative stable states. While we study dynamics in a setting where the fluctuations of OTUs of a same community can be considered independent, they are obviously not empirically. Moreover, both types of fluctuations are mediated by the same interaction between species and the environment. Understanding the structure of these correlation [11, 23] would be the first step to understand how the alternative stable states emerge from dynamics.

## III. MATERIALS AND METHODS

### A. Data

We analyze time series of 14 individual coming from three different datasets: ten individuals of the BIO-ML dataset [24] (all those for which a dense long-term time-series is available), the two individuals M3 and F4 from the Moving Pictures dataset [17] and the two individuals A and B from [3]. The length of the time-series ranges from 6 months to 1.5 year, and the sampling frequency varies (daily in the most dense series). Individuals A and B from [3] both undergo a period of strong disturbance to their gut flora due, respectively, to two diarrhoea episodes during a travel abroad and a *Salmonella* infection. We exclude these periods from the analysis and consider for each individuals two separate time-series, before and after the perturbation. More details on the data analysis can be found in the SI.

### B. Measuring dissimilarity

We want to measure the dissimilarity between the composition of a community at time *t* and at time *t* + *T*. Let 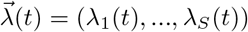 be the relative abundance of the S OTU in the community at time *t*. At time *t*, *N*(*t*) sequences are sampled, resulting in counts 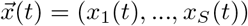. The relative OTU abundances 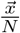 found in the two samples at lag *T* differ due to the difference in relative OTU abundances in the community, 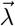, but also due to random sampling. In fact, even if we sampled twice the same community we would find some differences in the measured relative OTU abundances, especially if the sampling depth *N* is small with respect to the size of the community. Therefore, any dissimilarity measure computed directly on the measured relative abundnces 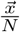 would overestimate the dissimilarity. The overestimation, additionally, is more pronounced for rare OTUs and for samples with smaller sampling efforts. To correct for the bias introduced by random sampling, we use a dissimilarity measure, Φ_i_(*T*), that is defined on the real relative abundance in the two communities, 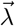, but can be estimated from the sampled counts 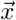. The dissimilarity for OTU *i* between time *t* and *t* + *T* is defined as

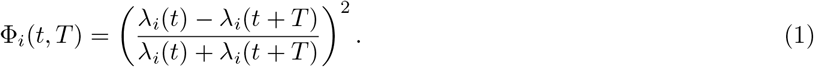

Φ_*i*_ takes values in [0,1]. It is equal to 0 when the relative abundances are equal and to 1 if the relative abundance is zero at one time and non-zero at the other. If we define *d_i_*(*t, T*) = *x_i_*(*T*) — *x_i_*(*t + T*) and *s_i_*(*t, T*) = *x_i_*(*T*) + *x_i_*(*t + T*), we can prove (see SI) that, if *N*(*t*) = *N*(*t + T*) (which can be obtained simply down-sampling the sample with larger *N*),

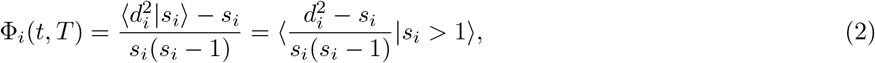

where 〈·〉 represents the average over realizations of the sampling, and is constrained to those realizations where *s_i_* > 1. This average over realization cannot be computed from the data, as we have a single realization. However, its average over time t is approximated by the average over time of the single realizations (SI).

Therefore, we compute the dissimilarity at lag T by averaging over the time-points of the time-series

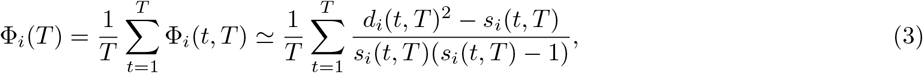

where now *d_i_*(*t,T*) and *s_i_*(*t,T*) are the values in a single realization of the sampling, known from the data. In this way, we obtain an estimation of the dissimilarity of real relative abundances *λ_i_* at a lag *T* based on the sampled counts *x_i_*.

### C. Estimating parameters of the dynamics from data

Previous studies [15, 16] showed that the dynamics of the relative abundance λ of a microbial species is best described by the stochastic logistic model with environmental noise

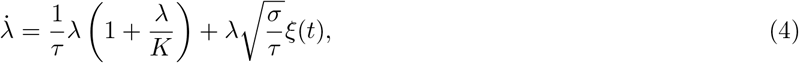

where *ξ*(*t*) is Gaussian white noise. This model has three parameters: *τ* has the dimension of a time, and determines the time-scale of relaxation to stationarity, *K* would be the carrying capacity in the absence of noise, and *σ* measures the intensity of the environmental noise. The model does not include interaction among species, and therefore cannot reproduce patterns of inter-species correlation, but correctly reproduces several patterns of the dynamics of a single species [15, 16]. Most importantly, if *σ* < 2 the stationary distribution is Gamma

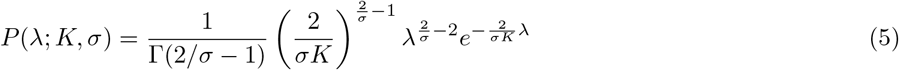

with mean 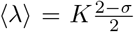 and variance 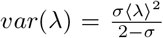. Note that the coefficient of variation 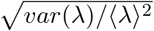 depends only on the parameter *σ*, which can thus be interpreted as the amplitude of the fluctuations.

The parameters *K* and *σ* can be estimated from the mean and variance of the abundance time-series. The variance of abundance fluctuations is also influenced by sampling. We use the sampling-corrected estimate as done in [16]

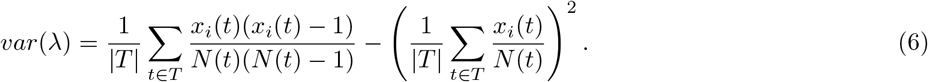

We note that the variance estimated with this formula may result negative if many counts are 0 or 1. The OTUs for which this happens are excluded from the analysis, as it is not possible to estimate their parameters.

### D. Theoretical expected values for dissimilarity

We can compute the expected value of the dissimilarity of an OTU between two snapshot of the same community at a large lag *T*, 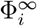, by assuming that the abundances at the two times are i.i.d. Gamma variables distributed according to (5) with the same parameters, equal to the estimated *K* and *σ* for that OTU. Note that this is the expected value of 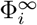 under the assumption that the dynamics is stationary. The expected value can be computed analytically, and it depends only on *σ*:

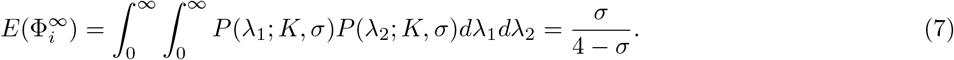

Similarly, we can compute the expected value of 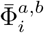, taking the two abundances to be Γ random variables with different parameters:

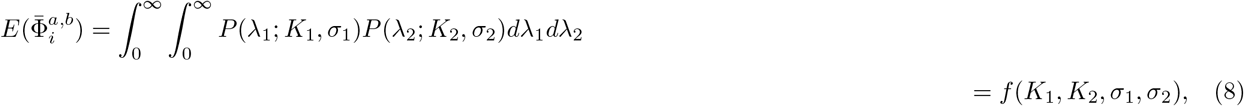

where 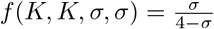 See the SI for the expression of *f*.

### E. Identifying OTUs with a plateau in intra-individual dissimilarity

To classify the curves 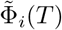 as either flat or increasing, we performed a linear fit discarding the initial transient (*T* > 10) and classified as increasing those with slopes above a threshold. To establish the threshold, we accounted for the fact that, although we expect a flat dissimilarity if the abundance is stationary, a slope different from zero can be find due to the noise in the estimation of Φ, which depends on the length and density of the time series, on the sequencing depth as well as from the OTUs parameters. Therefore, for each individual we computed the threshold as follows. We simulated the dynamics of each OTU according to the stochastic logistic model with parameters equal to the parameters estimated for that OTU and *τ* = 1. From these time series of 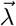 we sampled the time series of 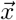 according to the sampling depth of the corresponding samples in the data, obtaining therefore values of 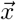 only for days for which the individual was sampled. We then computed 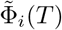 and its slope for each OTU. We defined as threshold for an individual the 95% quantile of the slopes obtained for that individual, i.e. the value such that only 5% of slopes obtained from the simulation are larger. Results are robust to variations of the thresholds (SI).

### F. Identifying jumps in the carrying capactity *K*

To identify the points of a time-series where the carrying capacity *K* has a jump, we first estimated *K* in window of length *w* = 50 days, so that for each day *t* we have two estimates, *K_for_*, estimated in the forward window (*t, t + w*], and *K_back_*, estimated in the backward window (*t — w, t*]. The estimates of *K* are computed using the estimate of *σ* for the entire time-series. Note that we accepted an estimate of *K* only if the corresponding window contains at least 5 samples. If the count are null for all samples in a window, the value of *K* is set to the value such that the probability to observe non-zero counts is 1/*w*.

The two estimates of *K* will differ on days around a change in *K*. To detect these days, we compute two quantities,

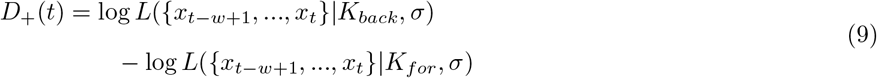

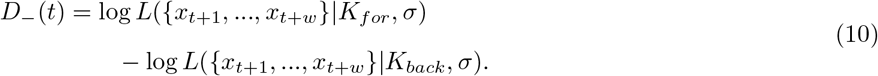

*D*_+_ has a peak when *K_for_* does not predict well the counts observed in the backward window, while *D*_ has a peak when *K_back_* does not predict well the counts observed in the forward window. Therefore, we identify the jumps in *K* as the times where *D*_+_ or *D*_ have a peak with height larger than a threshold (Fig S6). Results are robust to the variation of this threshold (see SI).

### G. Simulating abundance time-series with *K* jumps

We simulate abundance time-series according to a SLM with carrying capacity *K* that changes in time with discrete jumps. To simulate an individual, we consider the parameters *K* and *σ* of all its OTUs for which the parameters can be computed. For each individual, the values of 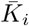 were set to the average of the estimated *K_i_* in all the individuals of the same dataset. Changes in *K* are a Poisson process, with rate such that the percentage of OTUs for which a jump is observed in the simulated time window is equal to the percentage observed empirically for that individual. The values of *K_i_* between jumps are extracted from 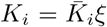 with a variance *ξ* of that is the same for all OTUs (equal to 2 for the results shown in Fig. 4, results are similar for different values, see Fig. S11). The obtained time-series are then sampled with a sampling depth equal to the average sampling depth for that individual, and only samples corresponding to days when the individual was sampled are kept (as the time-series density affects the noise in the dissimilarity). The obtained time-series of counts are analyzed as the empirical ones. Correlations are computed for the OTUs for which the parameters can be estimated.

