## Supplementary Information for "A macroecological description of alternative stable states reproduces intra- and inter-host variability of gut microbiome"

#### 1 Data

The data used in this study come from three different datasets: the Moving Picture (MP) dataset [1], the BIO-ML dataset [3] and the dataset from the study by David et al, called DA dataset in the following. [2]. Raw data for MP and D were obtained from MGnify [?], under project IDS MGYS00002184 and MGYS00001278, while for BIO-ML they were obtained from NCBI, under project ID PRJNA544527.

The MP dataset contains time-series of stool samples from two individuals, M3 (male) and F4 (female). M3 is sampled for $\sim 1$ year while F4 for $\sim 6$ months, with approximately daily sampling frequency. For the BIO-ML dataset, we considered the time-series of stool samples of 10 individuals (those with long and dense time-series): ae, am, an, ao, ba, bc, bh, bj, cl, co. The length of the time series varies from 6 months to 1.5 years, and the density varies (daily in the densest series). The DA dataset contains time-series of stool samples from two individuals, A and B. The two individuals are sampled over approximately 1 year with approximately daily sampling frequency.

Raw 16S sequences were processed with QIIME. The reads were initially processed with the split_libraries_fastq.py script with default parameters, then OTUs were picked using the script pick_closed_reference_otus.py, relying on UCLUST, with the Greengenes database at 97% similarity level as reference. We excluded samples with less than $10^4$ reads (13 samples in A and 6 samples in B), those with more than half sequences unassigned (one sample in A and one in B) and the samples identified as mislabelled in the original studies.

For individuals A and B from the DA dataset we consider two separate time series before and after the perturbation period (see Methods, section A): for individual A we consider the pre-travel (days 0 to 70) and post-travel (days 123 to 364) periods, and for individual B the pre-*Salmonella* (days 0 to 150) and post-*Salmonella* (days 160 to 252) periods. We note that, according to the original study [2], individual A after the perturbation period goes back to the pre-travel state, while individual B has a different composition post- *Salmonella*.

#### 2 OTU selection

For the analyses reported in Fig. 2 and Fig. 3, we used OTUs with average relative abundance $> 10^{-4}$. This choice is due to the fact that the estimation of $\Phi_i(T)$ is extremely noisy for rare OTU. Results are however similar for different thresholds (we tested $10^{-5}$, $10^{-3}$, $10^{-2}$).

For the analyses reported in Fig. 4 and 5, instead, we do not use this threshold as it would bias the correlation of parameters across-individuals. In fact, if we restricted ourselves to OTUs above a threshold abundance in both individuals we would bias towards OTUs that have a similar abundance in the two individuals. However, also keeping all OTUs for which we can estimate the parameters $K$ and $\sigma$ would bias the results. In fact, among the very rare OTUs we only observe those with high $\sigma$ (the others are either not observed or have too low counts to estimate $\sigma$ and $K$), see Fig. S1. Therefore, including OTUs that are too rare would yield a biased correlation of $\sigma$ across-individuals, as the rare OTUs that are observed in both individuals tend to have high values

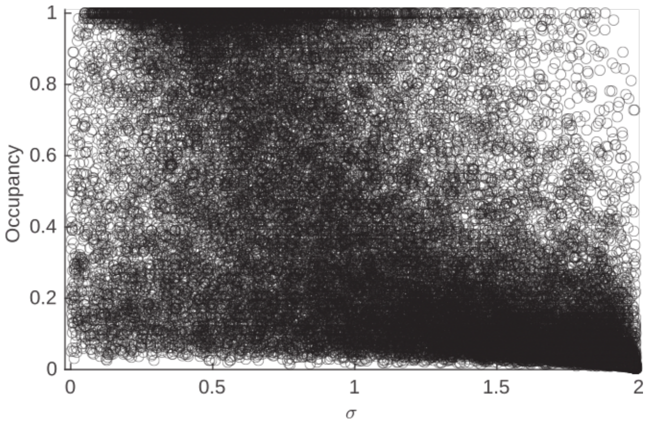


Figure S1: Observed OTUs with low occupancy (fraction of samples where they are observed) are biased towards high $\sigma$.

of this parameter. Based on the scatter plot of $\sigma$ and occupancy in Fig. S1, we therefore keep for the analysis all OTUs with occupancy $> 0.2$ (and for which it is possible to estimate $\sigma$ and $K$).

### 3 Estimate of $\Phi_i$ from sampled counts

We want to obtain an estimate of the dissimilarity

$$\Phi_i(t,T) = \left(\frac{\lambda_i(t) - \lambda_i(t+T)}{\lambda_i(t) + \lambda_i(t+T)}\right)^2 \tag{1}$$

from sampled counts $x_i(t)$ and $x_i(t+T)$. The counts $x_i(t)$ are a binomial random variable with success probability $\lambda_i(t)$ and number of extractions $N(t)$. When $N(t) \gg 1$ and $\lambda_i(t) \ll 1$, they can be approximated by a Poisson random variable with rate $\lambda_i(t)N(t)$.
If we define $d_i(t,T) = x_i(t) - x_i(t+T)$ and $s_i(t,T) = x_i(t) + x_i(t+T)$, we can prove (see below) that the average of $d_i^2$ over different realisations of the sampling constrained to a fixed $s_i$ is

$$\langle d_i^2 | s_i \rangle = s_i(s_i - 1)\left(\frac{\lambda_i(t)N(t) - \lambda_i(t+T)N(t+T)}{\lambda_i(t)N(t) + \lambda_i(t+T)N(t+T)}\right)^2 + s_i. \tag{2}$$

If the number of sampled sequences are the same at the two times (which can be obtained simply downsampling the sample with larger $N$), we have

$$\langle d_i^2 | s_i \rangle = s_i(s_i - 1)\left(\frac{\lambda_i(t) - \lambda_i(t+T)}{\lambda_i(t) + \lambda_i(t+T)}\right)^2 + s_i, \tag{3}$$

that is, for $s_i > 1$,

$$\Phi_i(t,T) = \frac{\langle d_i^2 | s_i \rangle - s_i}{s_i(s_i - 1)} = \langle \frac{d_i^2 - s_i}{s_i(s_i - 1)} | s_i \rangle, \tag{4}$$

corresponding to Eq. (2) of the main text.

Let us now prove (2). Let $d = x_1 - x_2$, $s = x_1 + x_2$ where $x_1$ and $x_2$ are Poisson variables with rates $\mu_1$ and $\mu_2$.
The constrained moments of $d$ can be obtained from the derivatives of the constrained generating function

$$H_{d|s}(v) = \sum_d P(d|s)e^{vd} = \sum_d \frac{P(d,s)}{P(s)}e^{vd} = \frac{1}{P(s)}\sum_d P(d,s)e^{vd}. \tag{5}$$

$P(s)$ is simply the probability of the sum of two Poisson variables with rates $\mu_1$ and $\mu_2$, which is itself a Poisson variable with rate $\mu_1 + \mu_2$.

The quantity $\sum_d P(d,s)e^{vd}$ can be also computed explicitly using the moment generating function of $d$ and $s$:

$$\begin{aligned} H_{d,s}(v,u) &= \sum_{d,s} P(d,s)e^{vd}e^{us} = \sum_{d,s} P(x_1)P(x_2)\delta(x_1+x_2,d)\delta(x_1+x_2,s)e^{vd}e^{us} \\ &= \sum_{x_1,x_2} P(x_1)P(x_2)e^{v(x_1-x_2)}e^{u(x_1+x_2)} = \sum_{x_1} P(x_1)e^{v+u}\sum_{x_2} P(x_1)e^{u-v} \\ &= G_{\mu_1}(u+v)G_{\mu_2}(u-v), \end{aligned} \tag{6}$$

where $G_\mu(h)$ is the Poisson moment generating function. This can be computed easily:

$$G_\mu(h) = \sum_{x=0}^{\infty} P(x)e^{hx} = \sum_{x=0}^{\infty} \frac{\mu^x}{x!}e^{hx-\mu} = e^{-\mu}\sum_{x=0}^{\infty}\frac{(\mu e^h)^x}{x!} = e^{-\mu}e^{\mu e^h} = e^{\mu(e^h-1)} \tag{7}$$

Substituting this result in the expression for $H_{d,s}$ we get

$$\begin{aligned} H_{d,s}(v,u) &= e^{\mu_1(e^{u+v}-1)}e^{\mu_2(e^{u-v}-1)} = e^{e^u(\mu_1e^v+\mu_2e^{-v})-\mu_1-\mu_2} = e^{(\mu_1e^v+\mu_2e^{-v})(e^u-1)+\mu_1e^v+\mu_2e^{-v}-\mu_1-\mu_2} \\ &= G_{\mu_1e^v+\mu_2e^{-v}}(u)e^{\mu_1e^v+\mu_2e^{-v}-\mu_1-\mu_2}. \end{aligned} \tag{8}$$

Writing $H_{d,s}(u,v)$ in this form is useful because if we call $F^{-1}$ the inverse operation to taking the generating function, we have

$$\begin{aligned} \sum_d P(d,s)e^{vd} &= F_u^{-1}(\sum_{d,s} P(d,s)e^{vd}e^{us}) = F_u^{-1}(H_{d,s}(u,v)) = F_u^{-1}(G_{\mu_1e^v+\mu_2e^{-v}}(u)e^{\mu_1e^v+\mu_2e^{-v}-\mu_1-\mu_2}) \\ &= F_u^{-1}(G_{\mu_1e^v+\mu_2e^{-v}}(u))e^{\mu_1e^v+\mu_2e^{-v}-\mu_1-\mu_2} = \frac{(\mu_1e^v+\mu_2e^{-v})^s}{s!}e^{-\mu_1e^v-\mu_2e^{-v}}e^{\mu_1e^v+\mu_2e^{-v}-\mu_1-\mu_2} \\ &= \frac{(\mu_1e^v+\mu_2e^{-v})^s}{s!}e^{-\mu_1-\mu_2}. \end{aligned} \tag{9}$$

Substituting this result in Eq. (5) we finally obtain the expression for the conditional generating function

$$H_{d|s}(v) = \frac{s!}{(\mu_1+\mu_2)^s}e^{\mu_1+\mu_2}\frac{(\mu_1e^v+\mu_2e^{-v})^s}{s!}e^{-\mu_1-\mu_2} = \left(\frac{\mu_1e^v+\mu_2e^{-v}}{\mu_1+\mu_2}\right)^s \tag{10}$$

We can thus compute conditioned moments:

$$\langle d\rangle_s = \frac{dH}{dv}|_{v=0} = s\frac{\mu_1-\mu_2}{\mu_1+\mu_2}$$

$$\langle d^2\rangle_s = \frac{d^2H}{dv^2}|_{v=0} = s(s-1)\left(\frac{\mu_1-\mu_2}{\mu_1+\mu_2}\right)^2 + s\ ,$$

which corresponds to equation 2.

#### 4 The time average of the averages over realisations is approximated by time average of single realisations

Let $[\cdot]$ denote the average over time, i.e. $[X] := \frac{1}{T}\sum_t x_t$, and $\langle\cdot\rangle$ denote the average over realisations. We want to show that $[\langle X\rangle] \to [X]$ when $T \to \infty$. To do that, we can show that the generating function of the two variables are equal in that limit.
$[X]$ is a stochastic variable with generating function $H_{[X]}(h) = \sum_{[X]} P([X]|\{p_1, ..., p_T\})e^{h[X]}$. Instead, $[\langle X\rangle]$ is deterministic, but we can think of it as a stochastic variable delta distributed around its value, with generating function $F(h) = \sum_x \delta(x, [\langle X\rangle])e^{hx} = e^{h[\langle X\rangle]}$.

$$\begin{aligned} H_{[X]}(h) &= \sum_{[X]} P([X]|\{p\})e^{h[X]} = \sum_{[X]}\sum_{x_1,...,x_T} p(x_1, ..., x_T|\{p\})\delta([X], \frac{\sum x_i}{T})e^{h[X]} \\ &= \sum_{x_1,...,x_T} p(x_1, ..., x_T|\{p\})e^{h\frac{\sum x_i}{T}} = \sum_{x_1,...,x_T}\prod_{i=1}^{T} P(x_i|p_i)e^{h\frac{x_i}{T}} = \prod_{i=1}^{T}\sum_x P(x|p_i)e^{h\frac{x}{T}} \\ &= \prod_{i=1}^{T} G_{x|p_i}\left(\frac{h}{T}\right) = \prod_{i=1}^{T}\exp\left(\log\left(G_{x|p_i}\left(\frac{h}{T}\right)\right)\right) = \exp\left(\sum_{i=1}^{T}\log\left(G_{x|p_i}\left(\frac{h}{T}\right)\right)\right) \\ &\simeq \exp\left(\sum_{i=1}^{N}\frac{h}{T}\langle X\rangle\right) = e^{h[\langle X\rangle]}, \end{aligned} \tag{11}$$

where to get the last row we used the fact that $\log G_x\left(\frac{h}{T}\right)$ is the generating function of the cumulants of $x$, therefore when $T \gg 1$ we can expand it as $\frac{h}{T}\langle X\rangle + \frac{1}{2}\left(\frac{h}{T}\right)^2 var(X) + \ldots$ and we cut the expansion after the first term.

#### 5 Expected value of $\Phi_i^{a,b}$ under Gamma fluctuations

The expected value of the dissimilarity of OTU $i$ across two individuals $a$ and $b$ can be obtained by taking the expected value of

$$\Phi_i^{a,b}(t, T) = \left(\frac{\lambda_i^a(t) - \lambda_i^b(t+T)}{\lambda_i^a(t) + \lambda_i^b(t+T)}\right)^2, \tag{12}$$

where the relative abundances $\lambda_i^a$ and $\lambda_i^b$ are independent Gamma distributed random variables with different parameters, $(K_a, \sigma_a)$ and $(K_b, \sigma_b)$. The expected value can be derived analytically with

Wolfram Mathematica, and yields

$$
\begin{aligned}
E(\Phi_i^{a,b}) = f(K_1,K_2,\sigma_1,\sigma_2) = -\frac{1}{K_2^2(1-\sigma_2)\sigma_2^3}\Big[\sigma_2^2\Big(-K_1^2(\sigma_1-2)-2K_1K_2(\sigma_1-2)(\sigma_2-1) \\
+K_2^2(\sigma_2-2)(\sigma_2-1)\Big)+\frac{2}{\sigma_1^2}\left(\frac{K_2\sigma_2}{K_1\sigma_1}\right)^{2/\sigma_1}\Gamma\left(2\left(\frac{1}{\sigma_1}+\frac{1}{\sigma_1}-1\right)\right) \\
\times\Big(\sigma_1(\sigma_2-2)[-2K_1K_2(\sigma_1-2)(\sigma_1-\sigma_2)(\sigma_2-1)+K_1^2(\sigma_1-2)(\sigma_1\sigma_2-\sigma_1-\sigma_2) \\
+2K_2^2(\sigma_2-1)(\sigma_1\sigma_2-\sigma_1-\sigma_2)]_2F_1\left(\frac{2}{\sigma_1},\frac{2}{\sigma_1}+\frac{2}{\sigma_2}-1;\frac{2}{\sigma_1}+\frac{2}{\sigma_2};1-\frac{K_2\sigma_2}{K_1\sigma_1}\right)\frac{1}{\Gamma(2/\sigma_1+2/\sigma_2)} \\
-2\sigma_2(K_1^2(\sigma_1-2)((\sigma_1\sigma_2-\sigma_1-\sigma_2)+2K_1K_2(\sigma_1-2)-2+(\sigma_2-1) \\
+2K_2^2(\sigma_1+\sigma_1(\sigma_2-2)\sigma_2^2+\sigma_2(-3+(5-2\sigma_2)\sigma_2))) \\
\times_2F_1\left(\frac{2}{\sigma_1}+1,\frac{2}{\sigma_1}+\frac{2}{\sigma_2}-1;\frac{2}{\sigma_1}+\frac{2}{\sigma_2};1-\frac{K_2\sigma_2}{K_1\sigma_1}\right)\frac{1}{\Gamma(2/\sigma_1+2/\sigma_2)}\Big)\Big]\,,
\end{aligned}
\tag{13}
$$

where ${}_2F_1$ is the ordinary Hypergeometric function.

### 6 Results are robust to variation of the threshold slope used to identify increasing $\Phi_i(T)$

For the results shown in Figures 2C and D and 3C, we identified OTUs with increasing $\Phi_i(T)$ by using a threshold slope different for each individual (see Methods). Results remain qualitatively similar if we use a common threshold for all individuals, defined as the median of the individual thresholds, see Fig. S2. The main differences are seen for individuals A before travel and B after Salmonella, as they have an individual threshold that is much larger that the median threshold.

### 7 Results are robust to variation of the threshold value of $D_+$ and $D_-$ used to identify jumps of $K$

The threshold value of $D_+$ and $D_-$ ($D_{thres} = 40$) used in the main text to identify jumps of the carrying capacity $K$ was chosen based on a good trade-off between false-positive and false-negative identifications, as judged by eye-inspection. However, results are qualitatively similar if the threshold is varied between 30 and 70, see Fig. S3.

### 8 Variance of the carrying capacity variability $\xi$

We hypothesized that the carrying capacity of an OTU $i$ can be modelled as

$$K_i = \bar{K}_i\xi, \tag{14}$$

where $\bar{K}_i$ is an OTU-specific carrying capacity and $\xi$ is a lognormal random variable with mean 1 and variance that is the same for all OTUs, independent from $\bar{K}_i$. To verify this assumption we computed $\bar{K}_i$ and var($\xi$) for the OTUs that are present in all 10 individuals of BIO-ML. For each OTU, we obtained $\bar{K}_i$ as the average of the $K_i$ estimated in each individual. Then, inverting Eq. (14) we obtained 10 values of $\xi$ for each OTU, of which we computed the variance. The obtained variances are plotted in Fig. S4A against $\bar{K}_i$. Although the obtained variances display a pattern, with larger variances corresponding to intermediate carrying capacities, such pattern can be explained simply

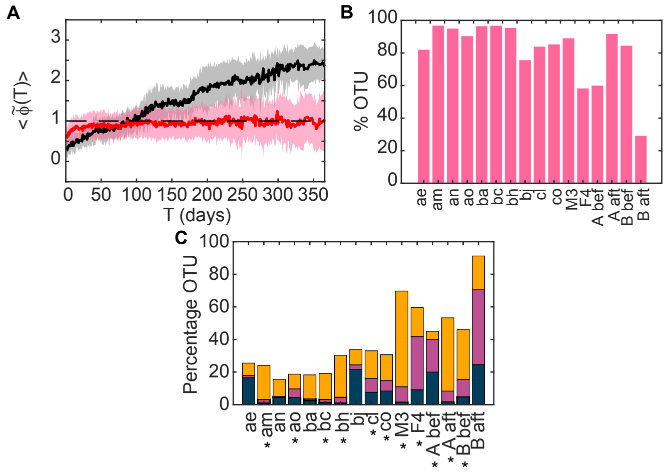


Figure S2: Robustness of results when the threshold slope used to identify increasing $\Phi_i(T)$ is the median of the individual thresholds used in the main text. A) Average of $\tilde{\Phi}(T)$ over OTUs with $\tilde{\Phi}$ classified as flat (red curve) and increasing (black curve), for individual 'bh'. Shaded areas represent one standard deviation intervals; B) Percentage of OTU whose $\tilde{\Phi}(T)$ is classified as flat in each individual; C) Percentage of OTUs where we detect a jump in $K$ and whose $\tilde{\Phi}$ is increasing (orange bars), where we detect a jump in $K$ but $\tilde{\Phi}$ is flat (yellow bars) and where $\tilde{\Phi}$ is increasing but we detect no jump in $K$ (blue bars).

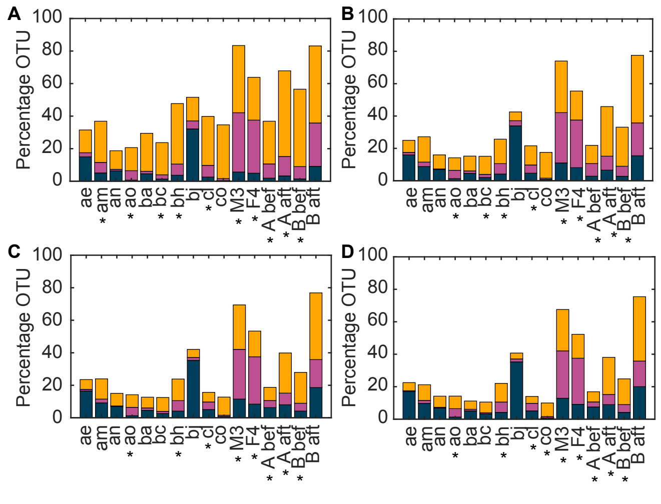


Figure S3: Robustness of results to variation of the threshold used to identify peak of $D_+$ and $D_-$. Panels A-D: Percentage of OTUs where we detect a jump in $K$ and whose $\tilde{\Phi}$ is increasing (orange bars), where we detect a jump in $K$ but $\tilde{\Phi}$ is flat (yellow bars) and where $\tilde{\Phi}$ is increasing but we detect no jump in $K$ (blue bars), for four different value of the threshold: 30, 50, 60, 70.

by an observation bias. In fact, OTUs with low $\bar{K}_i$ are observed in all 10 individuals only if var($\xi$) is sufficiently low, otherwise the $K_i$ in some individuals will fall under the observation threshold. On the other hand, when $\bar{K}_i$ is large the variance is bounded as $K_i < 1$. In fact, we can reproduce the empirical pattern from carrying capacities extracted from Eq. (14) with constant var($\xi$). We extract 10000 values of $\bar{K}_i$ from a lognormal distribution, and for each of these we compute 10 values of $K_i$ using var($\xi$)=5. We mimic observation by keeping only the OTUs such that $K_i > 10^{-5}$ in all 10 individuals. On these, we perform the same analysis performed on the empirical data. The results, in Fig. S4B, show a pattern similar to the one found in empirical data. Therefore, we conclude that the values of $K_i$ observed in the data are compatible with the model in Eq. (14) with var($\xi$) independent of $\bar{K}_i$.

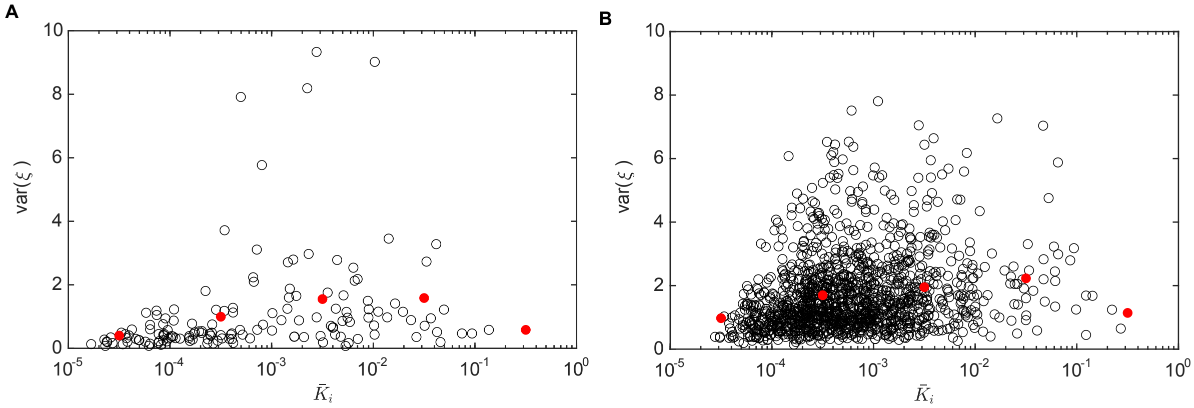


Figure S4: A) Variance of $\xi$ plotted against $\bar{K}_i$ for OTUs common to the 10 individuals of BIO-ML. Each circle is an OTU. Red circles are the average in bins of logarithmic size; B) Relationship between $\xi$ and $\bar{K}$ obtained from simulated data, according to Eq. (14).

### Supplementary figures and tables

| | | | |
|---|---|---|---|
| ae | 0.6439 | cl | 0.0000* |
| am | 0.0341* | co | 0.2162 |
| an | 0.8546 | M3 | 0.0001* |
| ao | 0.0000* | F4 | 0.0000* |
| ba | 0.2872 | A bef. travel | 0.0000* |
| bc | 0.0094* | A aft. travel | 0.0008* |
| bh | 0.0009* | B bef. Salm. | 0.0001* |
| bj | 0.7376 | B aft. Salm. | 0.9248 |

Table S1: P-values of hypergeometric test. Values with a * reject the null hypothesis that OTUs with transitions in the parameter $K$ are not over-represented among OTUs with increasing $\Phi(T)$.

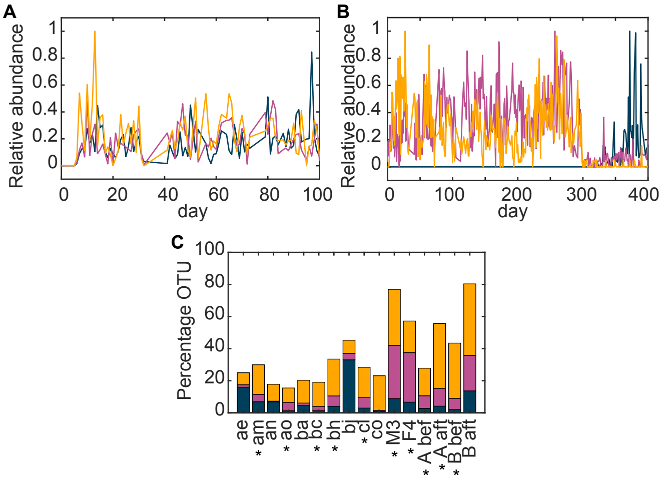


Figure S5: A and B: Examples of abundance time-series from individual 'M3' of the dataset Moving Pictures displaying sudden jumps in relative abundance, respectively, around day 5 and around day 300 (compare with corresponding peaks in Fig. S8); C) Percentage of OTUs where we detect a jump in $K$ and whose $\tilde{\Phi}$ is increasing (orange bars), where we detect a jump in $K$ but $\tilde{\Phi}$ is flat (yellow bars) and where $\tilde{\Phi}$ is increasing but we detect no jump in $K$ (blue bars).

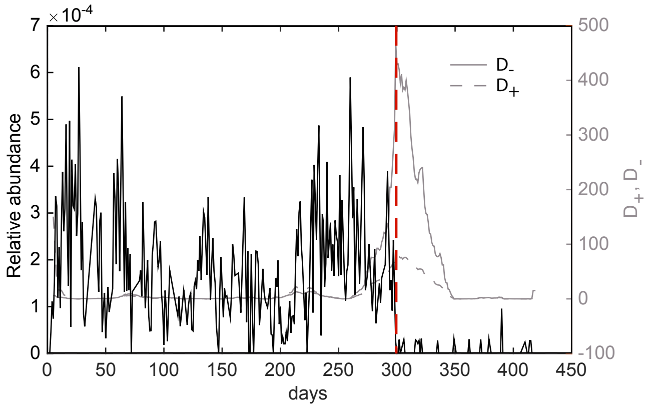


Figure S6: Example of our method to identify jumps in the abundance time series. A time series is plotted in black, and the corresponding $D_+$ and $D_-$ are plotted in grey. Both have a peak ($D > 40$) when the relative abundance has a sudden jump. The position of the peak (red dashed line) correctly identifies the jumping time.

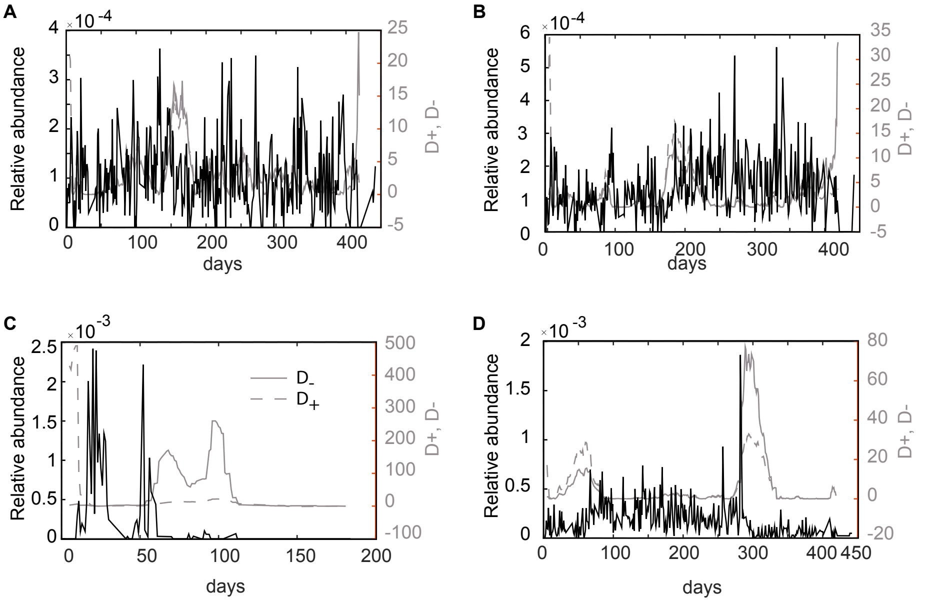


Figure S7: Examples of cases where $\Phi(T)$ is increasing even though no jump in $K$ is detected (A and B), or viceversa (C and D). A) No jump in relative abundance can be seen, $\Phi(T)$ has a positive slope due to noise; B) There is a jump in $K$ but the peaks of $D$ are lower than the threshold we set ($D = 40$); C) Two jumps in $K$ are detected, however $\Phi(T)$ is flat; D) One jump in $K$ is detected, however the change in relative abundance is small and does not yield a significant slope in $\Phi(T)$.

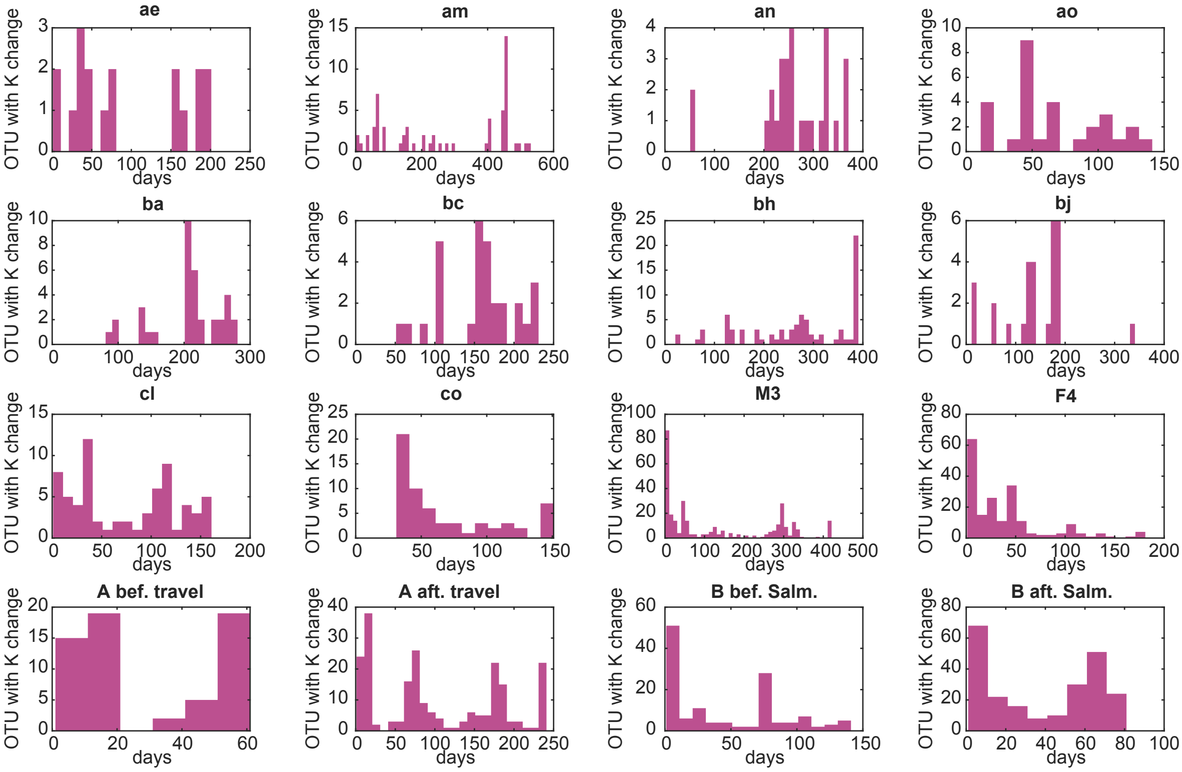


Figure S8: Histograms of times at which we detect a jump in the carrying capacity $K$ for all individuals.

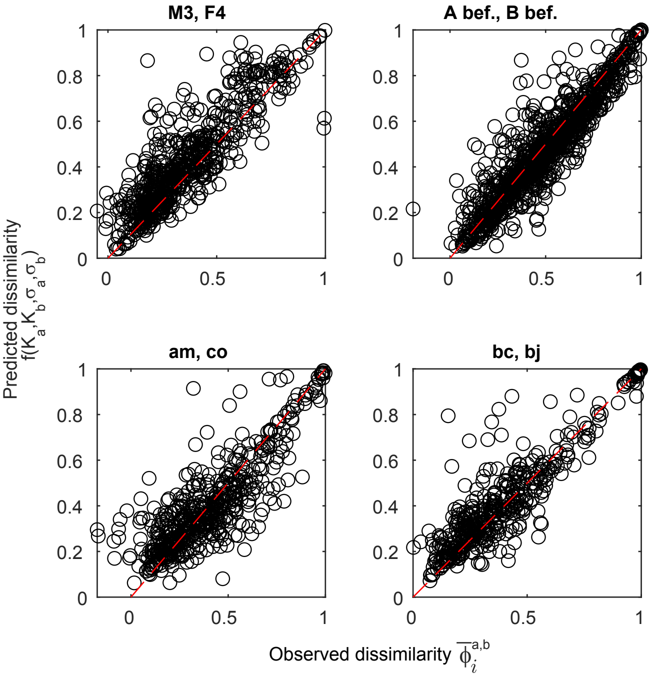


Figure S9: Comparison of the observed dissimilarity $\bar{\Phi}_i^{a,b}$ of OTU $i$ between a pair of individuals $a$ and $b$ with its theoretical expected value $f(K_a, K_b, \sigma_a, \sigma_b)$ computed with the individual parameters estimated for each individual. Each point represents an OTU. Pairs (M3, F4), (A bef., B bef), (am,co), (bc,bj) are shown, other pairs are similar.

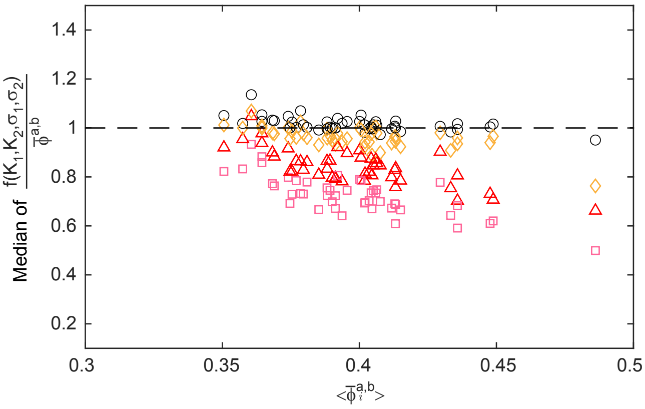


Figure S10: Median (over OTUs) of the ratio between the theoretical expectation for the dissimilarity of an OTU across two individuals, $f(K_1, K_2, \sigma_1, \sigma_2)$, and its empirical value $\bar{\Phi}_i^{a,b}$, plotted against $\bar{\Phi}_i^{a,b}$. The theoretical expected value captures typically all the observed dissimilarity $\bar{\Phi}_i^{a,b}$ when individual-specific parameters are considered in the prediction (black circles, $K_1 = K_a, K_2 = K_b, \sigma_1 = \sigma_a, \sigma_2 = \sigma_b$). The dissimilarity expected only from independent abundance fluctuations (pink squares, $K_1 = K_2 = \bar{K}, \sigma_1 = \sigma_2 = \bar{\sigma}$) accounts typically for roughly 70% of the total dissimilarity. The percentage however varies, being larger in individuals that are more similar. The remaining $\sim$ 30% of the empirical dissimilarity is mostly explained by the differences in the parameter $K$ across the two individuals (orange diamonds, $K_1 = K_a, K_2 = K_b, \sigma_1 = \sigma_2 = \bar{\sigma}$), while the differences in the parameter $\sigma$ has a smaller role (red triangles, $K_1 = K_2 = \bar{K}, \sigma_1 = \sigma_a, \sigma_2 = \sigma_b$ ). Each point corresponds to a pair of individuals from the same dataset.

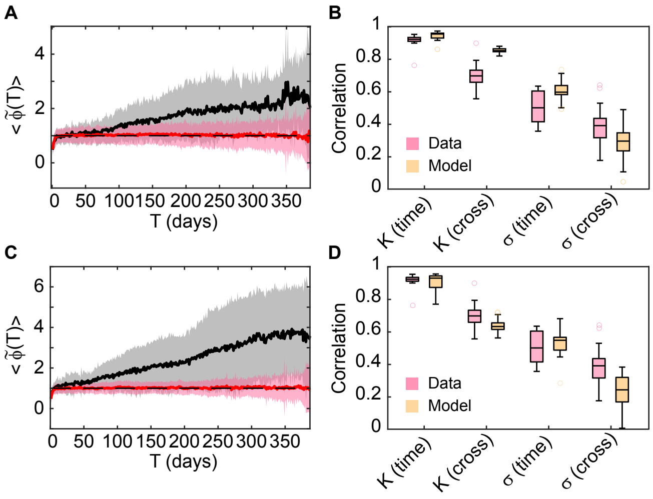


Figure S11: Comparison of model predictions with data, as in Fig. 5 of the main text, for two different values of $var(\xi)$: $var(\xi) = 1$ in panels A and B, $var(\xi) = 10$ in panels C and D.

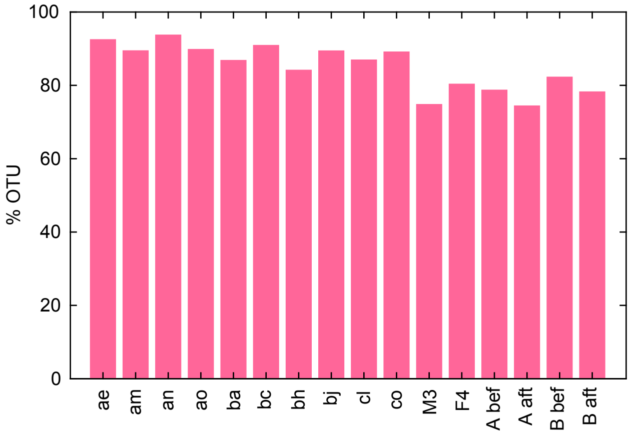


Figure S12: Percentage of OTU whose $\tilde{\Phi}(T)$ is classified as flat in each individual for simulated time-series (to compare with empirical results in Fig. 2D).

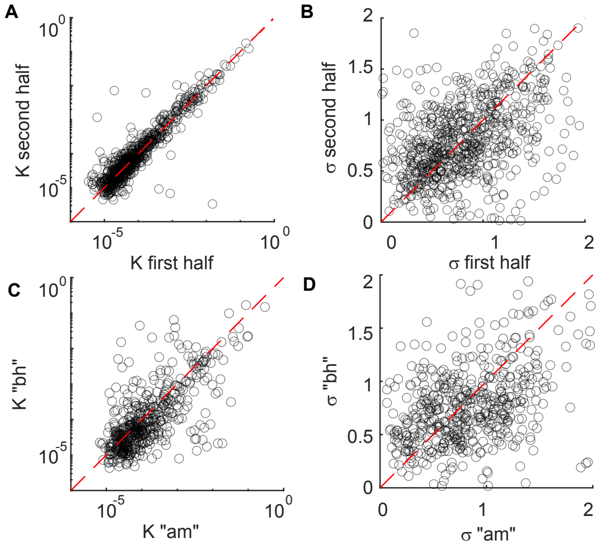


Figure S13: Example of consistency of parameters $K$ and $\sigma$ between two halves of a time-series and between two different individuals. A) and B): Scatter plots of, respectively, the carrying capacities $K$ (A) and the noise intensities $\sigma$ (B) estimated for each OTU on the first and second half of the time series of individual 'bh' from the dataset BIO-ML. Each point represents an OTU. The 1:1 line is shown as reference; C) and D): Scatter plots of, respectively, the carrying capacities $K$ (C) and the noise intensities $\sigma$ (D) estimated for each OTU in individuals 'am' and individual 'bh' from the dataset BIO-ML. Each point rep-resents an OTU. The 1:1 line is shown as reference.
